# Neural correlates of memory in a naturalistic spatiotemporal context

**DOI:** 10.1101/2022.11.30.518606

**Authors:** Matthew R. Dougherty, Woohyeuk Chang, Joseph H. Rudoler, Brandon S. Katerman, David J. Halpern, James P. Bruska, Nicholas B. Diamond, Michael J. Kahana

## Abstract

We investigated memory encoding and retrieval during a quasi-naturalistic spatial-episodic memory task in which subjects delivered items to landmarks in a desktop virtual environment and later recalled the delivered items. Transition probabilities and latencies revealed the spatial and temporal organization of memory. As subjects gained experience with the town, their improved spatial knowledge led to more efficient navigation and increased spatial organization during recall. Subjects who exhibited stronger spatial organization exhibited weaker temporal organization. Scalp-recorded electroencephalographic (EEG) signals revealed spectral correlates of successful encoding and retrieval. Increased theta power (*T* ^+^) and decreased alpha/beta power (*A*^−^) accompanied successful encoding, with the addition of increased gamma (*G*^+^) accompanying successful retrieval. Logistic-regression classifiers trained on spectral features reliably predicted mnemonic success in held-out sessions. Univariate and multivariate EEG analyses revealed a similar spectral *T* ^+^*A*^−^*G*^+^ of successful memory. These findings extend behavioral and neural signatures of successful encoding and retrieval to a naturalistic task in which learning occurs within a spatiotemporal context.

## Introduction

Episodic memory refers to our ability to remember events as occurring within a particular place and at a particular time (Tulving, 1972). Retrieval of an event tends to bring to mind others that occurred at nearby times (Kahana, 1996) and in nearby places (Miller et al., 2013), indicating that the spatiotemporal context of our experiences provides an organizational framework that supports later retrieval. Researchers have conducted investigations to better understand how we perform the feat of episodic memory using both classic list memory paradigms, such as free and cued recall (Murdock, 1974; Crowder, 1976; Kahana, 2012), and more naturalistic experimental approaches (Uitvlugt & Healey, 2019; Diamond & Levine, 2020). Measures of brain activity during such investigations have augmented purely behavioral memory measures, providing valuable insights into both episodic memory encoding and retrieval (Paller & Wagner, 2002; Johnson & Knight, 2015). However, owing to the technical difficulties of recording brain activity in naturalistic settings, this work has relied almost exclusively on experiments involving traditional list-memory tasks. Such experiments lack important features of everyday life, where experience flows continuously through time and space, and events occur through interaction with our environment.

The present paper addresses this knowledge gap by investigating the behavioral and electrophysiological correlates of encoding and retrieval in a quasi-naturalistic memory task. Specifically, we had subjects play the role of a courier, navigating through a virtual town on a bicycle and delivering items to target locations. Upon arrival at each location, subjects learned what item they delivered. On each ‘delivery day’ of this *Courier* task, subjects completed a route and delivered 15 unique items to unique businesses along the way. Then, after navigation to a 16th business, a blank screen prompted subjects to freely recall all of the delivered items from this delivery day. Following free recall, subjects completed a cued recall task, attempting to recall the item delivered at each (randomly) cued business location. A cumulative final free recall task followed each five delivery days. Across multiple sessions, subjects completed between 30 and 80 delivery days with the same subject-specific virtual town layout, allowing them to develop a mental representation of the town’s layout.

Having subjects encode and subsequently recall items delivered at variable times and in distinct spatial locations allowed us to utilize the order of recalls to evaluate the temporal and spatial organization of the encoded items. The free navigation aspect of the Courier game further allowed us to evaluate the role of spatial learning by measuring the degree to which subjects improved their navigational efficiency across repeated delivery days. Analyses of recall dynamics thus allowed us to ask several specific questions about the interacting roles of temporal and spatial information in memory retrieval. First, we sought to determine whether and to what degree subjects in the Courier task organize their recalls based on the temporal and spatial relations among the studied items (we examined these effects both on the immediate recall test following each delivery day and on the delayed cumulative recall test that followed each block of five delivery days). Previous findings in related tasks have found strong temporal clustering and a modest degree of spatial clustering (Miller, Weidemann, & Kahana, 2012; Pacheco & Verschure, 2018; Herweg, Sharan, et al., 2020). Observing spatial clustering effects in our task would validate the hypothesis that our paradigm encouraged subjects to encode items in relation to their spatial context. To the extent that these clustering effects arose, we additionally asked whether they were indicative of organizational efficiency at retrieval through analyses of inter-response times. Thirdly, we sought to understand the interplay between spatial and temporal organization of memories by evaluating the correlation between highly clustered recalls at various levels. Finally, to better understand the role of spatial context in organizing and influencing memories, we evaluated subjects’ ability to learn their spatial environment and the degree to which such proxies for learning influenced the organization of their recalls.

Along with the behavioral responses, we also recorded continuous high-density scalp EEG data. These data allowed us to evaluate several key hypotheses about the neural correlates of successful memory encoding and retrieval in the Courier task. Before describing our EEG hypotheses, we summarize key EEG correlates of successful encoding and retrieval identified in list-memory tasks. Numerous prior studies have documented a distinct pattern of spectral changes accompanying successful encoding and subsequent episodic recollection. In an early intracranial EEG study, Sederberg, Kahana, Howard, Donner, and Madsen (2003) documented what we refer to as the *T* ^+^*A*^−^*G*^+^ of successful memory. Analyzing individual electrodes, they found that increases in 4 Hz theta activity (*T* ^+^), decreases in 10 − 20 Hz alpha (and to some degree beta band activity (*A*^−^)), and increased > 40 Hz Gamma activity (*G*^+^) accompanied the encoding of subsequently recalled items. Subsequent intracranial studies have replicated the alpha decreases and gamma increases in large subject samples (Burke et al., 2014). Still, the theta increases have been somewhat elusive (Herweg, Solomon, & Kahana, 2020) except for studies that either specifically focused on the lowest frequencies within the theta band (Lin et al., 2017) or studies that separated narrowband and broadband components (Rudoler et al., 2022). Studies of episodic recollection have observed similar spectral patterns of retrieval success. For example, Burke, Merkow, Jacobs, Kahana, and Zaghloul (2015) reported theta increases in frontal regions, and widespread alpha decreases and gamma increases associated with the spontaneous recall of previously studied items. Similar spectral signatures of successful memory appeared across a range of tasks (Greenberg, Burke, Haque, Kahana, & Zaghloul, 2015; van Vugt, Schulze-Bonhage, Litt, Brandt, & Kahana, 2010; Osipova et al., 2006; Griffiths, Parish, et al., 2019; Griffiths, Mayhew, et al., 2019).

Scalp EEG studies of word list tasks have uncovered similar spectral signatures of successful encoding and recall. Sederberg et al. (2006) identified gamma power increases and low-frequency power decreases as markers of successful encoding. Notably, they identified a dissociation between these markers as a function of study timing, with gamma power increases associated with early list item encoding and low-frequency power decreases associated with late list item encoding. Long, Burke, and Kahana (2014) built upon this research in another word list free-recall task, demonstrating a more detailed *T* ^+^*A*^−^*G*^+^ pattern associated with successful item encoding. Katerman, Li, Pazdera, Keane, and Kahana (2022) found that spectral *T* ^+^*A*^−^*G*^+^ predicted successful free recall of items studied on prior sessions (after delays of > 18 hours), suggesting that *T* ^+^*A*^−^*G*^+^ is not limited to short-term memory procedures.

Here we ask whether similar *T* ^+^*A*^−^*G*^+^ patterns appear during successful memory encoding and retrieval in Courier, where learning occurs dynamically as subjects navigate through a rich synthetic environment. Ideally, patterns of brain activity mediating successful learning and recall in a word list task would generalize to more naturalistic settings. However, several distinctive aspects of word list tasks caution against this conclusion. First, strategic processes, including rehearsal (repeating items in one’s mind) and creating verbal or imagery-based mediators to link items, can lead to successful subsequent recall (Murdock, 1974; Kahana, 2012). These strategic processes surely have physiological correlates, and these would appear as biomarkers of successful learning. In Courier, however, following each encoding event, subjects faced the challenge of navigating through a visually complex environment to find the next business. This inter-item task relied upon subjects’ memory for business locations and their ability to efficiently search the environment for each business. Under these conditions, one might expect different learning strategies, potentially leading to an absence of some or all of the spectral *T* ^+^*A*^−^*G*^+^ of encoding. Indeed, some theorists have suggested that encoding strategies in a list recall task may not closely resemble encoding under more naturalistic conditions (Hintzman, 2016). Further, the engagement of spatial coding in our task might also lead to specific strategies, such as remembering an item in relation to its spatial location within the environment. Such spatial encoding processes could have distinct neural sequelae. Finally, the literature on the enactment effect (Nyberg, 1993) indicates superior memory for enacted events compared to memory for items. Encoding of delivered items is intrinsically tied to the action of delivering them to a given business in this task, as compared to static memory of word presentations in a word list task. If one theorizes that the *T* ^+^*A*^−^*G*^+^ of successful memory reflects strategic processes engaged during static item encoding, then the action of delivering encoded items may modify the degree to which *T* ^+^*A*^−^*G*^+^ is seen. Recording EEG across multiple sessions allowed us to further evaluate whether machine-learning approaches that have successfully decoded mnemonic success in tasks with discrete, sequentially encoded items also decode performance in a task in which subjects experience items as they navigate ad-lib through a quasi-naturalistic task environment.

The field of cognitive neuroscience is rapidly evolving, with technological advances to neuroimaging systems pointing towards a future of research far less limited by the confines of today. Bridging the gap between the theoretical foundations set forth in rigidly-defined experimental conditions and the exploratory work to come with advancing technology (e.g., with mobile EEG recordings) is sure to prove difficult. The primary motivation of our neural analyses is to provide a much-needed theoretical steppingstone. As previously described, even with the same recording systems, lab conditions, and recall paradigms, the differences between word list and quasi-naturalistic tasks produce a wide variety of differences in hypothetical outcomes, with little background to support any one outcome over another. Identifying the neural correlates of successful memory in a quasi-naturalistic task environment will help to provide a hypothetical standing ground for future naturalistic batteries to come.

## Methods

### Subjects

As we sought to detect EEG correlates of successful memory encoding and retrieval in this task, we turned to predicate studies in the literature for guidance in determining sample size. The most closely related paper examining EEG correlates of memory encoding in free recall was published by Sederberg et al. (2006). They asked 35 subjects each to study and freely recall word lists, collecting data on 48 lists of 15 items for a total of 1,680 encoding lists. Their study revealed significant EEG correlates of successful memory encoding, and as such, it provided a meaningful benchmark for our study’s sample size. As another benchmark, Katerman et al. (2022) examined EEG correlates of successful retrieval in free recall in 40 subjects who each provided five sessions of recall. Using these studies as guidance, we set out to achieve at least the same level of statistical power here.

Over a ten month period, fifty young adults recruited among the students and staff at the University of Pennsylvania and neighboring institutions consented to the aforementioned study. Recruitment materials informed interested individuals of the video game-like nature of the study and cautioned individuals with a high propensity for motion sickness. Nine subjects dropped due to scheduling issues, and eleven subjects dropped due to feelings of motion sickness. The thirty remaining young adults (ages 18–27, 13 Females) each contributed between three and eight sessions of navigation and recall data. The data collected from this experiment acted as training data for a separate experiment. As such, we encouraged subjects to partake in as many sessions as their schedules allowed (up to eight). A selection of subjects’ schedules only allowed for a limited set of sessions, resulting in the range of three to eight sessions. The final data comprised of 1,840 lists across 30 subjects, achieving the desired level of statistical power set out by prior related literature (Sederberg et al., 2006; Katerman et al., 2022).

### Courier hybrid spatial-episodic memory task

#### Virtual Town

The virtual town consisted of 17 businesses and 15 non-identified buildings (see Figure 1C for an overhead map of the town). The town also contained a park and smaller props such as trees, benches, trash cans, and mailboxes. All businesses were visually distinct, with various unique features (including large banners displaying the businesses’ names) identifiable from a distance (see Figure 1A for an example street view). Town layout remained consistent across subjects. However, target location banners varied randomly across subjects (e.g. the location of the Craft Store for one subject could be the location of the Bakery for another). For a given subject, the locations of businesses remained consistent across all sessions. The environment was created and displayed using an in-house C# library that interacted with Unity, a widely used game engine. We created the 3-D models used in the virtual environment using the Town Constructor 3 Asset Pack on the Unity Asset Store.

**Figure 1.**
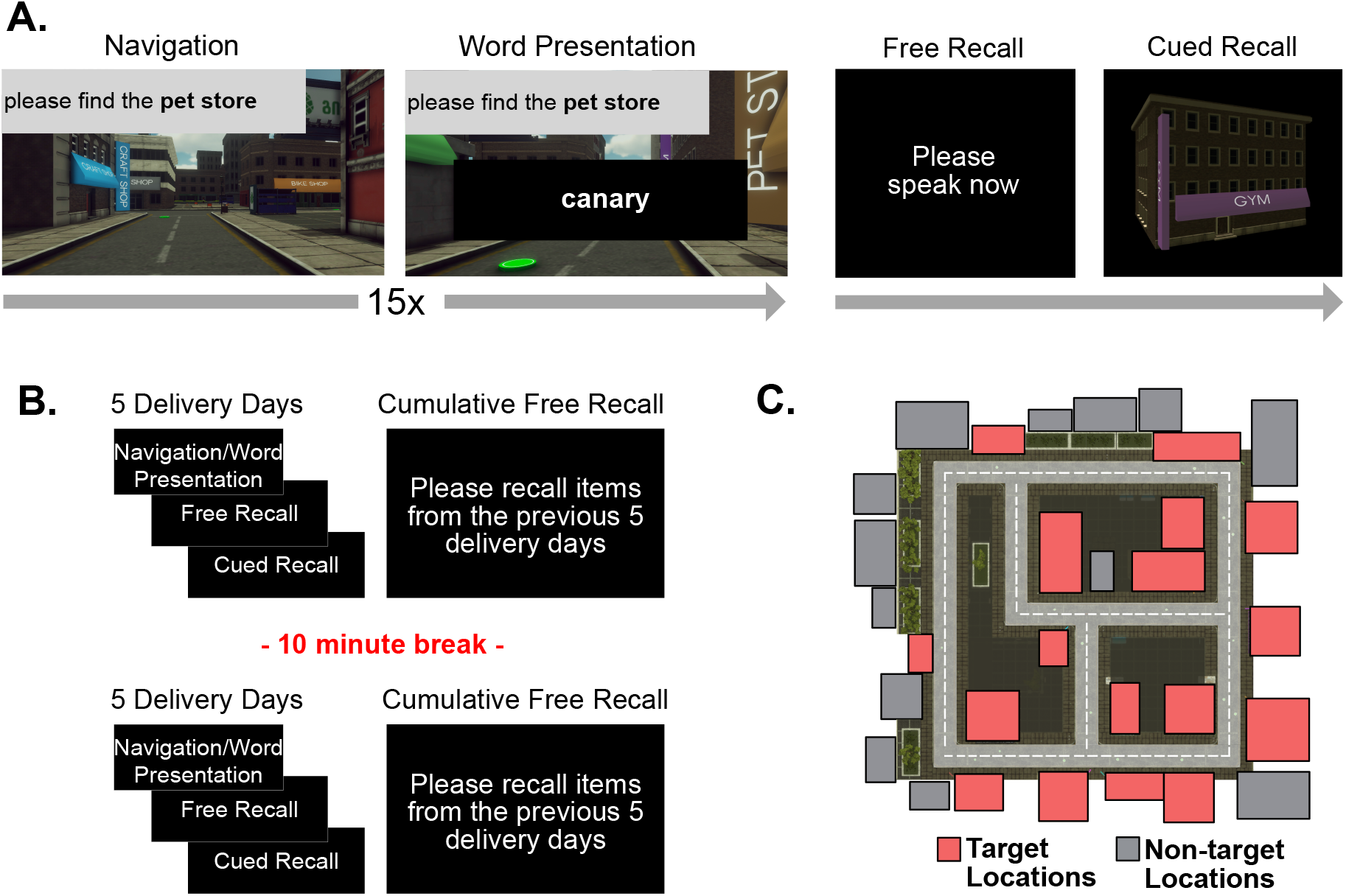
Task Design. **A**. A single trial within the Courier task. Subjects navigate to a specified business to deliver an item. Upon arrival, a word appears on screen indicating the item that was delivered. Subjects then navigate to the following business. After completing 15 deliveries, subjects complete free recall and cued recall of the delivered items. **B**. Structure of one session of the Courier task. Subjects complete five delivery days, followed by cumulative free recall of all items delivered across preceding delivery days. After a short break, subjects complete an additional five delivery days and cumulative free recall of items from these five delivery days. **C**. Top-down view of the virtual town. The town consists of 17 target locations, i.e. businesses, and 13 non-target locations.

#### Game Play

Subjects played the role of a courier in a hybrid spatial-episodic memory task, riding a bicycle and delivering parcels to businesses located within a virtual town using a video game controller. The experiment played comparably to desktop and console navigation-based video games. Each experimental session consisted of a series of delivery days (trials) during which subjects navigated to deliver items and subsequently recalled these items (See Figure 1A). Before starting the first delivery day of the first session, subjects participated in a “town learning” task. During this time, subjects navigated between each of the 17 target locations according to on-screen instructions generated in pseudorandom order without delivering items. Subjects navigated using the joystick and buttons on a video game controller. In the town learning phase and in each delivery day, an arrow dynamically pointing in the direction of the target location appeared on the street after 12 seconds of navigating to ensure the experiment ran within a reasonable window of time.

Each delivery day consisted of a navigation phase followed by two recall tasks (see Figure 1A). For the navigation phase of each delivery day, 16 of the 17 total businesses were chosen as target locations in a pseudo-random sequence subject to the constraint that business *i* + 1 could not be seen from the location of business *i*. This methodology ensured that subjects had to search for the next business guided by their knowledge of the spatial layout of the town. The randomization further minimized any positive correlation between spatial and temporal proximity. Subjects navigated to each location sequentially according to instructions presented on screen (see Figure 1A). Upon arrival at the first 15 businesses, subjects waited for 1000ms ± a 250 ms jitter, after which a word appeared on screen for 1600ms seconds while it was dictated aloud. These words, aka delivered items, never repeated within a session. Each delivered item was semantically related to its target business, however, items within a delivery day bore no particular semantic relation to one another. Immediately following word presentation, the next on-screen navigation instruction appeared (e.g. “please find the Pet Store”). At the 16th location, rather than item presentation, subjects saw a black screen and heard a beep, indicating the beginning of the free recall phase. During free recall, subjects had 90 seconds to recall as many items from this delivery day as they could in any order. Following completion of the free recall task, subjects completed a cued recall task. One business at a time appeared on screen for six seconds, during which subjects attempted to recall the item they delivered at this location. Vocal responses were recorded and annotated offline. Each session consisted of two identical blocks, each consisting of five delivery days and a two-minute cumulative free recall of the preceding five delivery days (see Figure 1B). A break of approximately ten minutes separated these two blocks. Sessions varied in timing based on the efficiency of navigation, with an average duration of roughly 90 minutes. The first session was a special case, in which the first block of trials prior to the break consisted of the town learning phase, one practice delivery day, two test delivery days, and cumulative free recall.

### Data collection and pre-processing

We recorded EEG with a 128-channel BioSemi ActiveTwo system with a 2048 Hz sampling rate. We applied a 0.1 Hz high-pass filter to remove baseline EEG drift over the course of each session. We then partitioned the recording into two segments, separating the period before and after the mid-session break. For each segment, we identified bad channels as those with excessive (|*z*| > 3) log-variance or Hurst exponent relative to other channels. We identified bad channels separately for each segment, as we reapplied gel to reduce impedance during mid-session breaks.^1^ We then dropped bad channels from their respective partitions and applied a common average reference scheme. We then reconstructed the original channels from the cleaned components, interpolated bad channels using spherical splines, and applied two fourth-order Butterworth notch filters, one with a 58-62 Hz stop-band and one with a 118-122 Hz stop-band, to remove electrical line noise.

### Data and Code Availability

All raw data and code for this study are freely available. Behavioral and electrophysiological data may be downloaded from OpenNeuro (Markiewicz et al., 2021) at https://openneuro.org/datasets/ds004706. The public repository Kahana (2023) contains all experimental data and analysis code, as well as additional scripts for data handling and preprocessing.

## Behavioral Analyses

### Temporal and Spatial Clustering

We assessed temporal and spatial clustering in two ways. First, we calculated conditional probability of recall transitions as a function of temporal or spatial separation of delivered items (i.e., lag in presentation order between two items for temporal, and euclidean distance between delivery locations of two items for spatial). We tallied the actual and possible distances of possible transitions from a given recall to compute these conditional response probabilities (Kahana, 1996; Miller et al., 2012). Second, we followed the procedure of Polyn, Norman, and Kahana (2009) to calculate temporal and spatial clustering scores for each subject. To compute clustering scores, we calculated percentile rankings of each possible transition ordered according to their distance from the just recalled item, where 0% corresponded to the shortest distance and 100% to the farthest. The temporal and spatial clustering scores for a given transition were defined as the distance rank of the actual transition relative to all the possible transitions. We calculated each subject’s overall clustering scores by averaging the scores of all recalls across all sessions. High temporal/spatial clustering scores indicate recall of items close in time/space. This approach allowed us to relate a given subject’s overall reliance on temporal or spatial context to other relevant variables.

Because spatial distances and serial positions were not fully independent during encoding, we tested hypotheses about clustering scores using permutation tests. For each trial, we shuffled the order of recalled items and recomputed clustering scores according to the above method. We then averaged scores across subjects and repeated the process 1000 times to attain a null distribution of clustering scores. By calculating the proportion of the null distribution with higher clustering scores than the observed average, we were able to determine whether average clustering scores across all subjects were above chance.

### Excess Path Length Analyses

To find the shortest navigation path between businesses, we used the Breadth-First Search (BFS) algorithm, a well-known algorithm for calculating the shortest path between any two nodes in a graph. In order to apply the BFS algorithm, we first vectorized the town layout into a binary format (e.g., 0 for road and 1 for non-road items), where each point in the vectorized array became a “node”. For a given navigation period (i.e. navigation from business *A* to *B*), we computed the shortest route and calculated the total distance of this route by summing the euclidean distances of its nodes. We then calculated the excess path length by subtracting the shortest navigation distance from the actual navigation distance (our approach builds on similar approaches to measured excess path length in Newman et al. (2007) and Manning, Lew, Li, Sekuler, and Kahana (2014)).

## EEG Analyses

### Spectral decomposition

We applied a Morlet Wavelet transform (five cycles) to compute spectral power at each electrode for eight log-transformed frequencies ranging from 3-128 Hz, corresponding to 3 Hz, 5.13Hz, 8.77Hz, 14.99Hz, 25.62Hz, 43.90Hz, 74.88Hz, and 128 Hz. We then log-transformed the resulting signal and z-scored the logtransformed signal at each frequency and electrode pair to estimate deviation from mean power.

#### Epoch construction

We categorized encoding and retrieval events into two groups for comparison. Subsequent memory of encoding events during immediate free recall determined categorization of encoding events. During encoding, words appeared on screen for 1600ms. For encoding analyses, each wavelet was calculated between 300ms and 1300ms following presentation onset, with a 833.33ms mirrored buffer applied to both sides of the data. For retrieval analyses, we identified 1000ms preceding correct recall events during immediate free recall (i.e. first instance of each item’s recall from the just-completed delivery day), excluding events for which subjects were not silent during these 1000ms.

We compared successful retrieval epochs to matched “deliberation” periods. We replicated the methodology of Solomon et al. (2019) to identify deliberation periods for each successful retrieval event. First, for each successful retrieval event, we searched for a 1000ms interval during immediate free recall in a separate delivery day of the same session during which no vocalization took place. We prioritized deliberation periods that occurred within the same 1000ms of a recall window as the retrieval event, and if no match existed, we extended the interval within 2000ms surrounding vocalization onset of the retrieval event. We excluded all successful retrieval events without matched deliberation periods from analysis. As a result, each successful retrieval event was matched with one deliberation interval from a different list. The mean number of retrieval events per session across subjects was 101.36 (standard deviation = 18.38), and the mean number of matched deliberation events per session across subjects was 67.40 (standard deviation = 16.53). The number of successful retrieval events was roughly comparable to the number of successful encoding events. Each wavelet for successful retrieval events was calculated between 900ms and 50ms prior to vocalization onset, with a 833.33ms mirrored buffer applied to both sides of the data. In accordance with this method, we calculated each wavelet for matched deliberation events between 100ms and 950ms of the 1000ms window with a 833.33ms mirrored buffer applied to both sides of the data.

#### Regions of interest

We collapsed spectral power estimates across subsets of electrodes to generate average spectral patterns at eight frequencies × eight regions of interest (ROIs). In order to maximize spatial coverage while minimizing inter-region overlap, scalp EEG researchers traditionally define eight scalp EEG ROIs based on segmentation of the coronal, sagittal, and horizontal axes (Curran, 2000). Recent studies have applied this same methodology using similar EEG caps to those used in this study (Long & Kahana, 2017; Weidemann, Mollison, & Kahana, 2009; Katerman et al., 2022), and have yielded a variety of spatially-resolved neural findings. Most recently, Katerman et al. (2022) demonstrated pronounced high-frequency power increases during successful retrieval only in posterior regions of the brain, as well as theta power increases only in anterior-inferior regions. As such, we selected ROIs in accordance with this standard. Figure 6A provides a diagram of the ROIs utilized in neural analyses.

#### Multivariate Classifier Training

We trained L2penalized logistic regression classifiers based on item encoding events, and separately on item retrieval events during free recall. For each epoch corresponding to an encoded item, we trained 1024 electrophysiological features (128 channels × 8 log-spaced frequencies from 3-128 Hz) against binary labels reflecting memory success. For encoding, this label indicates successful or failed retrieval during subsequent free recall. For retrieval, this label distinguishes correctly recalled words from periods of silent deliberation. We re-sampled the training data to ensure balanced class weighting. To select the L2 penalty parameter, we performed a nested leave-one-session-out cross-validation procedure, selecting the penalty term for the outer fold which achieved the highest average accuracy score across the inner folds.

To ensure proper model fitting, we only included subjects who recorded more than three sessions of training data, which reduced our sample size for this analysis to 29 subjects. Using less data would increase the risk of over-fitting the penalty parameter, especially in the inner folds of our nested cross-validation procedure.

#### Evaluating classifiers

We evaluated the accuracy of classifiers by computing the area under the curve (AUC) of the Receiver Operating Characteristic (ROC) function, which plots the proportion of true positives against the proportion of false positives at varying decision thresholds. We then asked how likely it was that this performance score would have been matched by a naïve classifier assigning labels randomly. In expectation, the AUC for random guessing is 0.5. However, we could not compare the observed within-subject AUC values to this theoretical null; since AUC scores can be highly correlated across cross validation folds (as the models shared most of their training data), so standard parametric tests would be biased in their estimate of the variance of within-subject model performance. Instead, we established a performance baseline via non-parametric permutation testing (Nichols & Holmes, 2001). Specifically, we randomly shuffled the true target labels and recomputed the classifier’s performance to generate an empirical null distribution of AUC values. An observed within-subject AUC greater than 95% of this null distribution met our standard for above-chance predictive accuracy.

To determine if predictive accuracy across subjects is significantly greater than chance, we computed a one-sample *t*-test on the distribution of observed AUC values against the expected mean of 0.5. This across-subject test is valid because subjects are independent.

To estimate the contribution of each frequency × ROI pair in the input space to classification performance, we constructed forward models based on the learned weights of the individual classifiers (Haufe et al., 2014). Raw feature weights in a backward model (like logistic regression) represent an extraction filter that is difficult to directly interpret. The weights express how the latent variable (here, mnemonic strength) can be decoded from input features. The weights do not necessarily express how the features vary as a function of the latent variable. This is especially true in a regularized regression setting, where weights are penalized to satisfy an additional term (the L2 norm) in the loss function. Feature activation values, however, actually express a generative model for the observed neural data as a function of the latent mnemonic status. For example, a positive feature activation for a theta-band feature can be interpreted as an indication that the model predicts theta power to be greater for subsequently recalled words.

## Results

Our data revealed several novel phenomena regarding both behavior and electrophysiology in the Courier task. We first report a detailed view of behavioral data, including analyses of recall initiation, the effects of serial position on recall probability, and the interacting roles of temporal and spatial clustering. We next describe how subjects learned about the spatial layout of the environment and the relation between the development of spatial knowledge and the organization of episodic memory. We then report on the spectral correlates of successful memory encoding and retrieval, testing the hypothesis that spectral *T* ^+^*A*^−^*G*^+^ effects previously reported in discrete item list-memory tasks emerge in our more quasi-naturalistic memory task. Next, we take a multivariate approach to our EEG data, asking whether we can reliably use spectral features to classify encoding and retrieval success and whether classifiers learn feature weights that resemble those seen in our univariate spectral analyses. Although secondary to our main research questions, we close by presenting behavioral results from our cued and cumulative free recall tasks.

### Recall dynamics in immediate free recall task

To analyze recall data, we treated each sequentially delivered item within a delivery day analogous to items presented in a typical list-memory task. This method created opportunities to evaluate recall performance as a function of serial position and to examine the influence of temporal and spatial relations between items on recall transitions. Figure 2A illustrates the serial position effect in our data. As each subject contributed data from 27-77 lists (across 3-8 sessions), we display individual subjects’ serial position curves (blue lines) along with the sample average (black line with 95% confidence intervals). Overall, subjects exhibited high levels of overall recall (*M* = 71.5%, *S D* = 12.1%); exceeding performance seen in traditional word list free recall tasks (e.g., Murdock, 1962; Brodie & Murdock, 1977). Although many procedural variables could have contributed to high recall rates, prior work suggests active encoding manipulations can enhance recall (Nyberg, 1993). Contextual variability may also have contributed to successful recall (Melton, 1970; Greene, 1989; Lohnas, Polyn, & Kahana, 2011).

**Figure 2.**
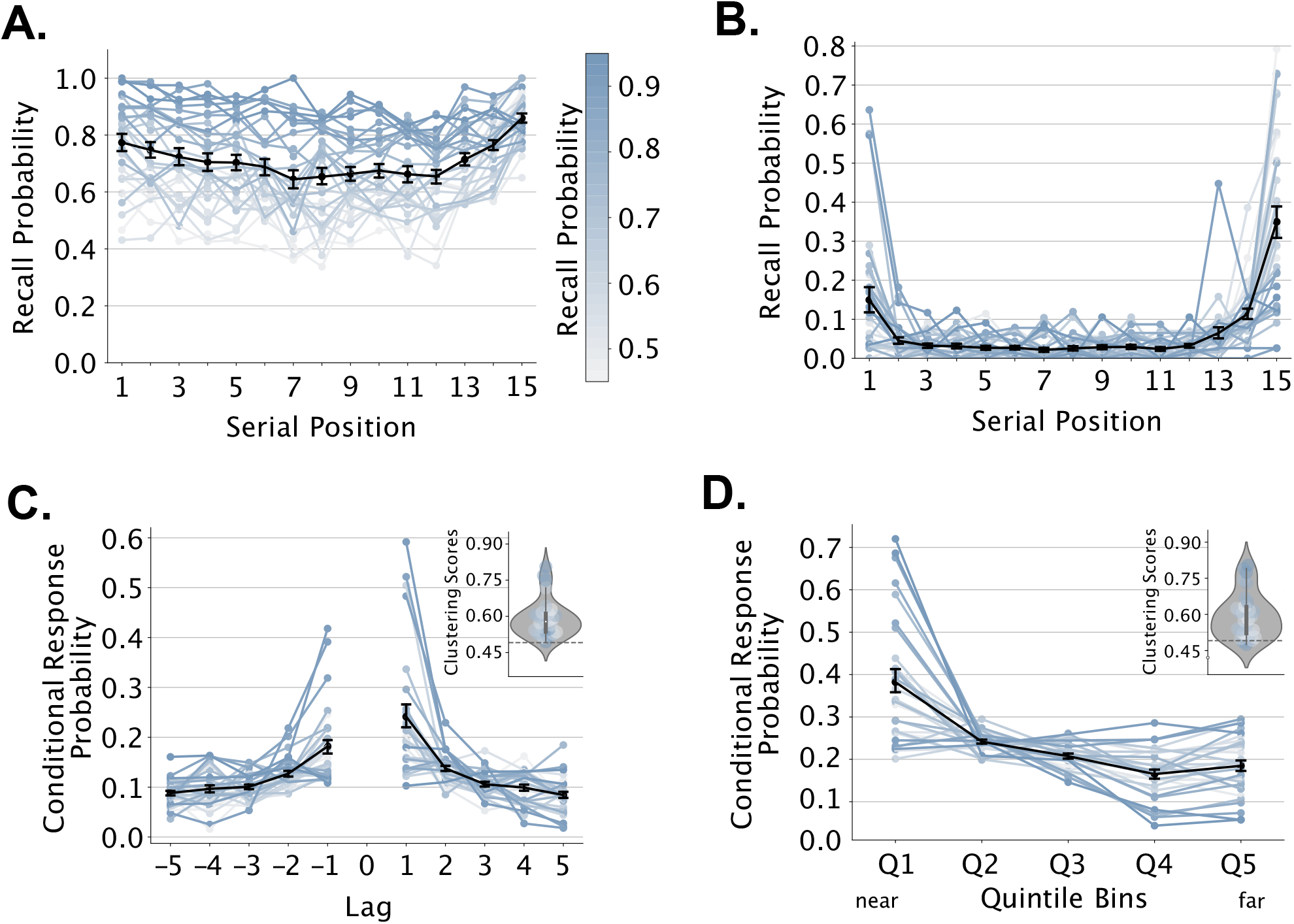
Free recall of delivered items. **A**. Probability of recalling delivered items as a function of their serial position (blue lines: individual subjects shaded by overall probability as indicated in color bar (applies to all panels); black line: sample average). **B**. Probability of initiating recall with a delivered item studied in a given serial position. **C**. The conditional response probability as a function of lag indicates the probability of successively recalling items from positions *i* and *i* + *lag* conditional on transition availability. Subplot shows the distribution of temporal clustering scores compared with chance clustering level (dotted line, see Methods). **D**. Spatial conditional response probability curve. Analogous to (C), we replace lag with Euclidean distance, binning possible distances between target locations into quintiles. Subplot shows subjects’ spatial clustering scores. Error bars in all panels indicate 95% confidence intervals, calculated using 1000 iterations of bootstrapped resampling of the data.

Subjects exhibited superior recall of primacy (serial positions 1-3 vs. 4-12, *t*_(29)_=5.64, *p* < 0.001, *d* = 0.52, 95% CI [0.05, 0.10]) and recency (serial positions 13-15 vs. 4-12, *t*_(29)_=4.80, *p* < 0.001, *d* = 0.92, 95% CI [0.06, 0.15]) items, though these effects appeared muted as compared with prior word list tasks (Murdock, 1962). Recall initiation showed a marked effect of serial position (Figure 2B), with subjects most often beginning recall with items from the end (serial positions 13-15 vs 4-12, *t*_(29)_=8.17, *p* < 0.001, *d* = 2.33, 95% CI [0.02, 0.08]), and beginning (serial positions 1-3 vs 4-12, *t*_(29)_=3.52, *p* < 0.001, *d* = 0.92, 95% CI [0.02, 0.08]) of a list. Individual subject graphs reveal a diversity of strategies, with some subjects consistently initiating their recall with the first list item and at least one subject initiating recall three items back. Prior studies with discrete word lists have documented all three of these strategies in similar proportions to those observed here (Healey & Kahana, 2014).

Although many prior studies have investigated recall dynamics in discrete word list tasks, few studies have considered dynamics of recall in more naturalistic tasks (e.g. Pacheco & Verschure, 2018). In Courier, an extended (and variable) period followed item delivery during which subjects searched for the next business. As such, subjects did not experience successively delivered items contiguously as they do in standard word-list free recall tasks. Nonetheless, we anticipated finding significant effects of temporal proximity, as these effects have been reported in related prior experiments (Miller et al., 2013) and in continual-distractor free recall tasks, where subjects perform a secondary distractor task between successively encoded words (Howard & Kahana, 1999). Although we also expected to observe a spatial clustering effect, earlier studies using similar methods have not always found reliable effects of spatial clustering (Herweg, Sharan, et al., 2020), or have reported clustering based on the shortest path separating locations rather than their distance in a metric (Euclidean) space. Observing spatial clustering using a Euclidean distance metric would suggest that subjects have formed an allocentric, map-like, representation of the environment (Manning et al., 2014).

Subjects exhibited a robust temporal clustering effect, successively recalling items delivered at proximate times during a given delivery day. In particular, the conditional probability of successively recalling deliveries *i* and *i* + *lag* falls off monotonically as |*lag*| increases (see Figure 2C; |lag| = 1 vs |lag| ≥ 4, *t*_(29)_=5.71, *p* < 0.001, *d* = 1.67, 95% CI [0.08, 0.16]). This temporal clustering effect also exhibits a strong forward asymmetry, indicating that subjects tend to make transitions from earlier to later deliveries, despite their overall tendency to begin recalling from the end of the list (Figure 2B; lag +1 vs −1, *t*_(29)_=4.47, *p* < 0.001, *d* = 0.56, 95% CI [0.03, 0.09])). Both of these effects appear consistently across subjects, but some subjects exhibit markedly stronger temporal clustering than others, as in word list tasks (Healey & Kahana, 2014).

As subjects experience each delivered item in a (virtual) location, we hypothesized that their recalls would exhibit a spatial clustering effect. For each recall transition, we evaluated spatial clustering by calculating the Euclidean distances between the just recalled item and all other possible recalls. Subjects’ tendency to successively recall items delivered to proximate spatial locations would signify a spatial clustering effect. Any measure of spatial clustering will make assumptions about how subjects encode spatial information. In what may be the first paper to examine spatial clustering in free recall, Miller et al. (2012) assumed that subjects represented spatial information as a graph. Other related work, however, has argued that individuals represent spatial information according to a Euclidean metric (O’Keefe & Nadel, 1978; Manning et al., 2014; Peer, Brunec, Newcombe, & Epstein, 2021). The latter assumption aligns with animal studies of place cells (neurons, typically in the hippocampus, that fire preferentially when an animal traverses a particular location), but assumes that subjects have a high level of familiarity with the environment, allowing them to transform their graph-like experience into a 2D spatial representation (Redish, 1999; Howard, Fotedar, Datey, & Hasselmo, 2005). In the present work, we followed Herweg, Sharan, et al. (2020) in assuming that subjects build a cognitive map representation in which the coordinates of businesses correspond to the spatial layout of the environment, without any distortion due to their experience. Prior work, however, has shown that experience can substantially distort the spatial map and that, in some cases, graph representations can be preferable (Warren, Rothman, Schnapp, & Ericson, 2017; Kuipers, Tecuci, & Stankiewicz, 2003). To the extent that subjects use these graph-like representations, they would reduce our measure of spatial clustering.

Subjects exhibited a strong spatial clustering effect (Figure 2D), validated through a repeated measures ANOVA and a post-hoc test comparing conditional recall probabilities of the nearest and farthest quintile bins (*F*(4, 116) = 27.88, *p* < 0.001,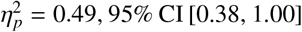*t*_(29)_=5.07, *p* < 0.001, *d* = 1.69, 95% CI [0.12, 0.28]). Similar to temporal clustering, subjects exhibited marked variability in their degree of spatial clustering, as illustrated in the subject-level spatial clustering curves, and the distribution of spatial-clustering scores, shown in the figure.

Given the significant influence of spatial and temporal similarity on subjects’ recall organization, we sought to better understand how these forms of organization relate to one another. We also wanted to confirm the existence of both forms of organization by ruling out the possibility that a hidden confound between temporal lag and spatial distance might produce an effect of spatial clustering. Although we designed Courier to minimize confounds between time and space (see *Methods*) if successively encountered businesses tend to occur in spatially proximate locations, a strong temporal clustering effect could produce an artifactual spatial clustering effect. We can test for the presence of this confound by evaluating the correlation between temporal and spatial clustering – a negative correlation between these forms of clustering accompanied by positive overall clustering effects, argues against the confound described above. Calculating clustering scores according to the aforementioned methods, we evaluated this relationship first across item recalls within delivery days and subsequently across subjects, following the work of Miller et al. (2012) and Herweg, Sharan, et al. (2020), respectively.

Figure 4 features the scatterplots and observed correlations between spatial and temporal clustering at both the delivery day and subject levels. We observed a significant negative correlation between these clustering scores at both the delivery day (*r*_(1838)_= − 0.28, *p* < 0.001, 95% CI [-0.31, -0.22]) and the subject (*r*_(27)_= − 0.73, *p* < 0.001, 95% CI [-0.80, -0.34]) levels, building upon findings in previous spatiotemporal episodic memory tasks (Herweg, Sharan, et al., 2020; Miller et al., 2012). These results provide evidence that subjects utilize distinct contextual cues during retrieval. Additionally, subject-level correlation findings suggest a consistency in subjects’ organization of recalls according to either spatial or temporal context, potentially indicative of conscious deployment of specific retrieval strategies.

Given that subjects demonstrated a tendency across the experiment to more regularly spatially or temporally cluster recalls, we further asked whether this tendency predicted their overall recall performance. Post-hoc correlation analyses of each subjects’ clustering scores with their overall recall probability across all sessions demonstrated a significant positive correlation between spatial clustering score and recall probability (*r*_(27)_=0.39, *p* < 0.05, 95% CI [0.01, 0.64]). Temporal clustering, however, only exhibited a very weak and non-statistically significant relation with recall probability (*r*_(27)_=0.14, *n*.*s*., 95% CI [-0.23, 0.47]). While the temporal relation of items at encoding varied throughout the experiment, the spatial information in the town layout remained consistent across trials and sessions. As a result, it is possible that subjects’ memory benefited from learning the environmental structure and using this knowledge to organize their recalls.

Following the method of Murdock and Okada (1970), we also analyzed inter-response times (IRTs) as a joint function of output position and of the number of items recalled. As shown in Figure 3A, IRTs grew exponentially with increasing output position, and did so more rapidly when subjects recalled fewer items in total. We next asked whether IRTs would similarly reveal effects of temporal and spatial clustering. Analogous to Figure 2C, we asked whether subjects exhibited faster transitions between successively recalled items experienced at proximate times during a given delivery day. Figure 3B shows that IRTs generally decreased with lag in both the forward and backward direction ( |lag| = 1 vs |lag| ≥ 4, *t*_(29)_= −6.47, *p* < 0.001, *d* = −1.14, 95% CI [ -1.82, -0.96]), but did not exhibit any evidence of asymmetry (lag +1 vs − 1, *t*_(29)_=0.54, *n*.*s*., *d* = 0.0, 95% CI [0.0, 0.0]). Applying the same logic to spatial clustering in Figure 3C, we observed faster transitions among successively recalled items experienced at nearby virtual locations (repeated measures ANOVA, *F*(4, 116) = 10.25, *p* < 0.001,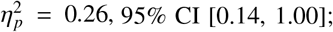post-hoc test Q1 vs. Q5, *t*_(29)_=−5.02, *p* < 0.001, *d* = −0.94, 95% CI [-1.83, -0.78]).

**Figure 3.**
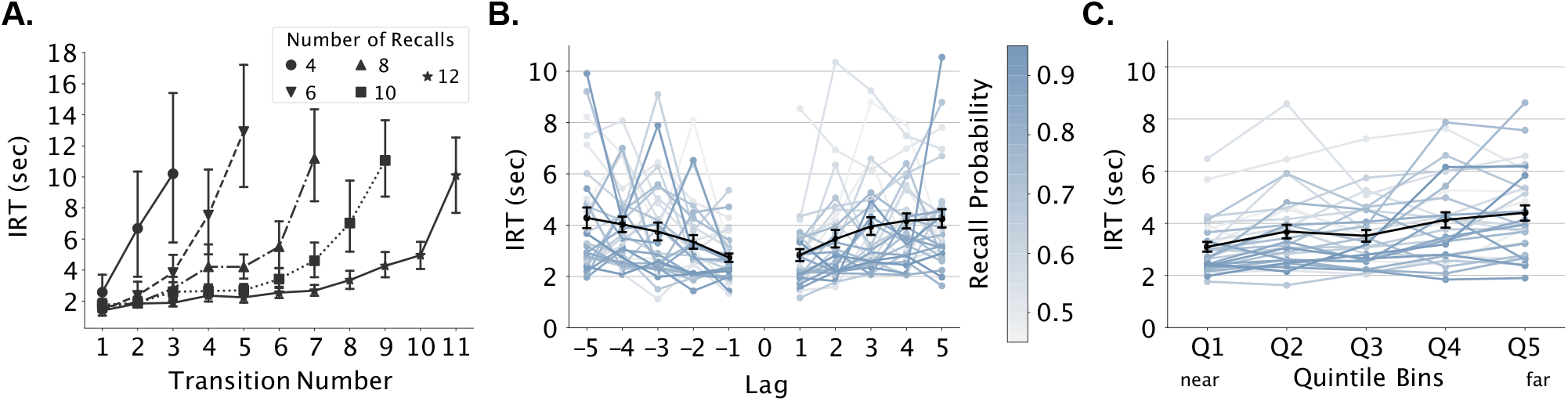
Inter Response Times (IRTs). **A**. IRTs for correct recall transitions increase as a function of output position (the numerical order of the retrieved items, e.g. one for the first retrieved item) and decrease as a function of total number of correct recalls. Transition number refers to the numerical order of transitions made during a recall sequence, e.g. one for the transition between items in the the first and second output positions. For clarity, we only show data for trials with an even number of correct recall transitions ranging from 4 to 12 (see legend). **B**. Conditional response latency as a function of temporal lag (blue lines: individual subjects shaded by overall probability as indicated in color bar (applies to all panels); black line: sample average) **C**. Conditional response latency curves as a function of distance bin. Analogous to (B), we replace lag with Euclidean distance, binning possible distances between target locations into quintiles. Error bars in all panels indicate 95% confidence intervals, calculated using 1000 iterations of bootstrapped resampling of the data.

**Figure 4.**
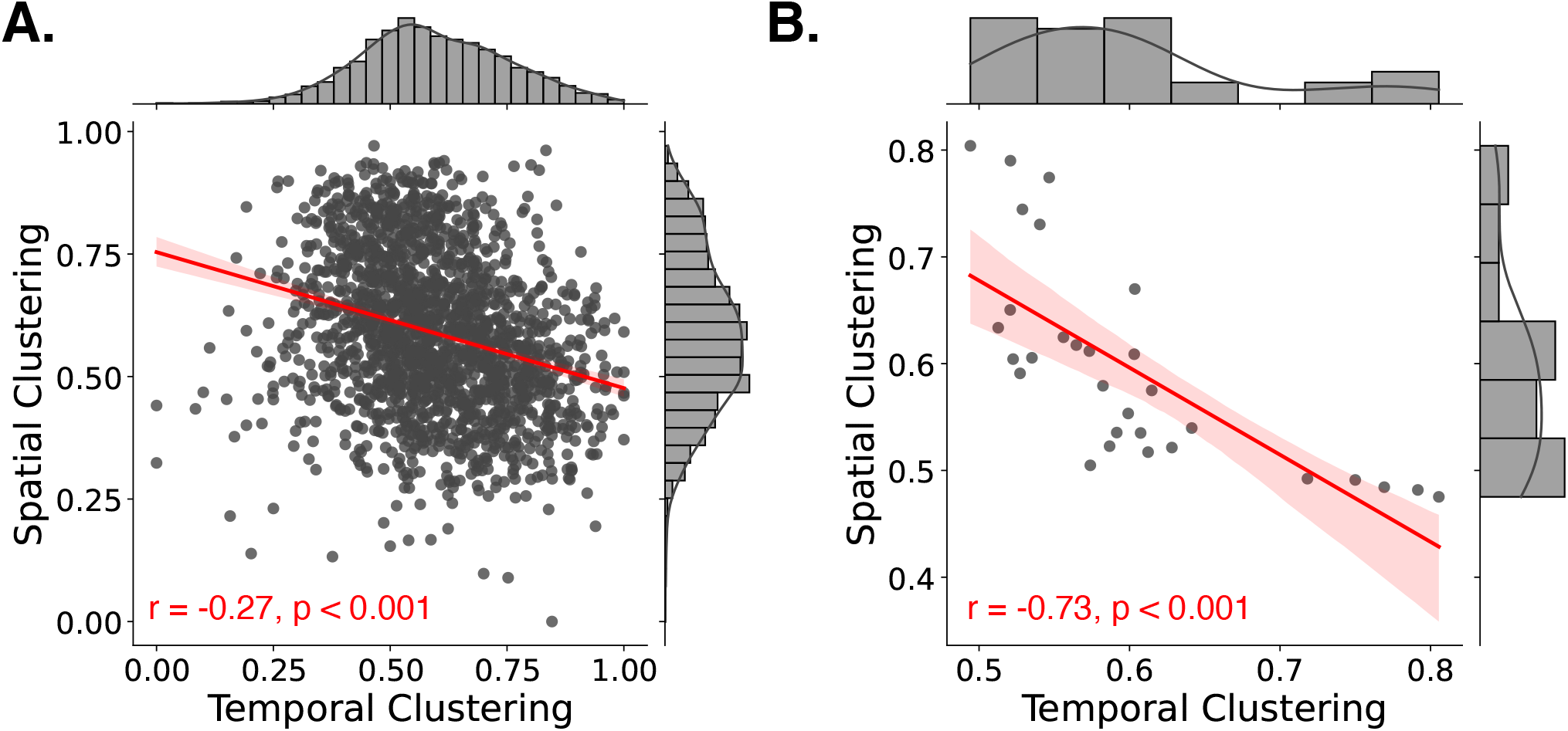
Correlation between spatial and temporal clustering. **A**. Correlation between spatial and temporal clustering scores of successful recalls for each delivery day. Each dot represents one delivery day. **B**. Correlation between subjects’ spatial and temporal clustering scores across all delivery days and sessions. Each dot represents one subject.

The spatial context of item delivery in our Courier task is unique compared to the temporal context in that subjects have the opportunity to build a mental representation of the town layout over time that can influence the organization of their memories. We confirmed that subjects learn their spatial environment across sessions through an analysis of excess path length, where short excess path lengths indicate town layout learning via increased path efficiency (see *Methods*). As shown in Figure 5B, excess path decreased as sessions progressed, indicating increased knowledge of the spatial environment with time (Manning et al., 2014) (repeated measures ANOVA, *F*(3, 78) = 16.24, *p* < 0.001, 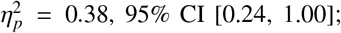post-hoc session one vs. four, *t*_(26)_=4.53, *p* < 0.001, *d* = 0.95, 95% CI [16.14, 42.04]). In addition, subjects demonstrated increased clustering of memories according to spatial context as sessions progressed (Figure 5C, repeated measures ANOVA, *F*(3, 78) = 4.22, *p* < 0.01,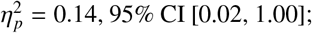post-hoc session one vs. four, *t*_(26)_= − 2.96, *p* < 0.01, *d* = −0.58, 95% CI [-0.09, -0.02]). More notable than subjects’ learning of their spatial environment is the relationship between this learning and the organization of their memories. In fact, subjects with shorter average excess path lengths demonstrated higher degrees of spatial clustering (*r*_(26)_= −0.42, *p* < 0.05, 95% CI [-0.67, -0.02], Figure 5D). Whereas temporal context provides a framework for organizing memories largely independent of learning and experience, the degree to which spatial context frames the organization of memory builds with experience. Given that the majority of our lives occur within a relatively consistent set of locations, these results suggest that the inclusion of spatial context in memory paradigms is valuablein developing an understanding of real-life mechanisms of memory organization.

**Figure 5.**
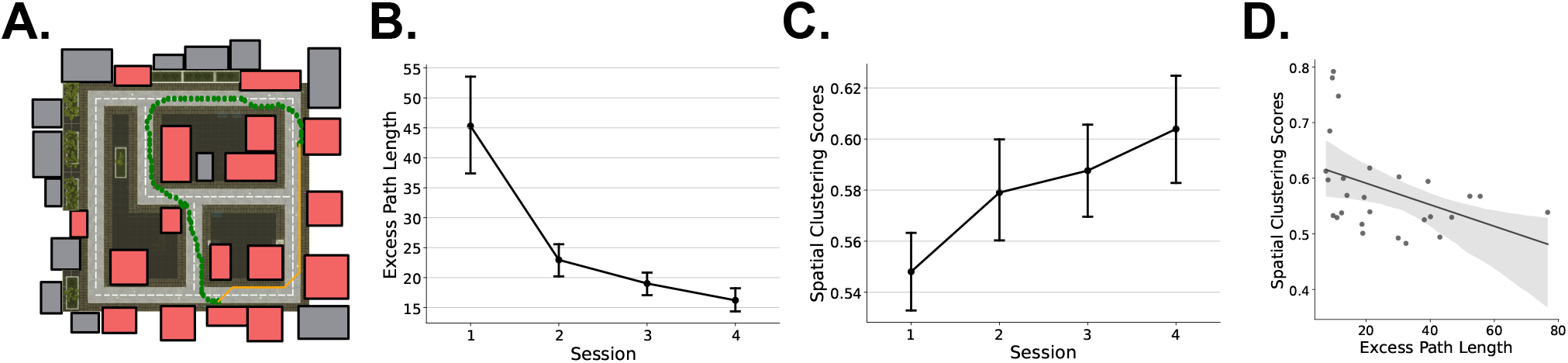
Environment Learning. **A**. Sample trace of actual navigation and shortest navigation paths. Green dots represent the actual navigation path an example subject took. The orange line represents the shortest path for that given navigation. See *Methods* for details on calculating the shortest path. **B**. Excess path length analysis. Excess path length is the difference between the shortest path distance and actual navigation path distance. **C**. Average spatial clustering scores as a function of session progression across subjects and sessions. **D**. Correlation between excess path lengths and spatial clustering scores. Each dot represents one subject’s data. For panels B, C, and D, we included data from the first four completed sessions in subjects who completed these sessions (*n* = 27). Error bars in panels B and C indicate 95% confidence intervals, calculated using 1000 iterations of bootstrapped resampling of the data.

### Spectral correlates of successful memory encoding and retrieval

Next, we sought to characterize EEG correlates of successful memory encoding and retrieval in our Courier task. Previous work in traditional word list tasks has identified increased theta power, decreased alpha/beta power, and increased gamma power as indicators of successful memory retrieval (Katerman et al. (2022) refer to this as a *T* ^+^*A*^−^*G*^+^ of successful memory retrieval). Studies of memory encoding have revealed similar findings, with particularly robust alpha/beta decreases (Long et al., 2014; Sederberg et al., 2006). Each of these results, however, have emerged in traditional discrete item list-memory tasks. Between the quasi-naturalistic aspects of our task, the robust evidence that subjects encode spatial information and use that information to guide search, and the disruption of memory processes during navigation and location search, this task provides a unique setting in which to investigate the generalizability of spectral *T* ^+^*A*^−^*G*^+^ effects outside of discrete list-memory tasks. Such generalizability gains theoretical relevance in light of arguments that word list recall tasks rely on encoding strategies that differ markedly from those used in our daily lives (Hintzman, 2016). To evaluate spectral correlates of memory encoding and retrieval in Courier we transformed the multivariate EEG time series into spectral power estimates between 3 and 128 Hz frequencies (see *Methods*). We then extracted mean power estimates during 1000ms item encoding epochs (between 300ms and 1300ms of the 1600ms total item presentation), and during an 850ms window preceding correct retrievals (900ms to 50ms prior to vocal onset, see *Methods*). All subsequent analyses used these spectral estimates.

We first consider the spectral correlates of successful memory encoding, comparing spectral estimates for the encoding of subsequently remembered and forgotten items. To do so, we first conducted an independent-samples t-test across events separately for each subject. This yielded *t*scores for each combination of electrode and frequency. Following this, we averaged the resulting *t*-scores across electrodes within an ROI to control for residual noise from any particular electrode. Figure 6B averages these mean *t*-scores across subjects to determine a variance-normalized effect size for each ROI × frequency pair. Here we see striking decreases in alpha/beta power during the encoding of subsequently remembered items. We also see increases in low-theta power during successful memory encoding. A clear pattern did not emerge in the gamma band. To assess the reliability of these effects across subjects, we conducted a one-sample *t*-test comparing the resulting average ROI × frequency *t*-scores to zero. Each ROI × frequency pair that met a *p* < 0.05 FDR-corrected criterion appears outlined in black. All ROIs, except for anterior-inferior regions, exhibited statistically reliable alpha/beta band decreases. Statistically significant theta increases appeared in left anterior inferior, left anterior superior, and right parietal superior regions.

**Figure 6.**
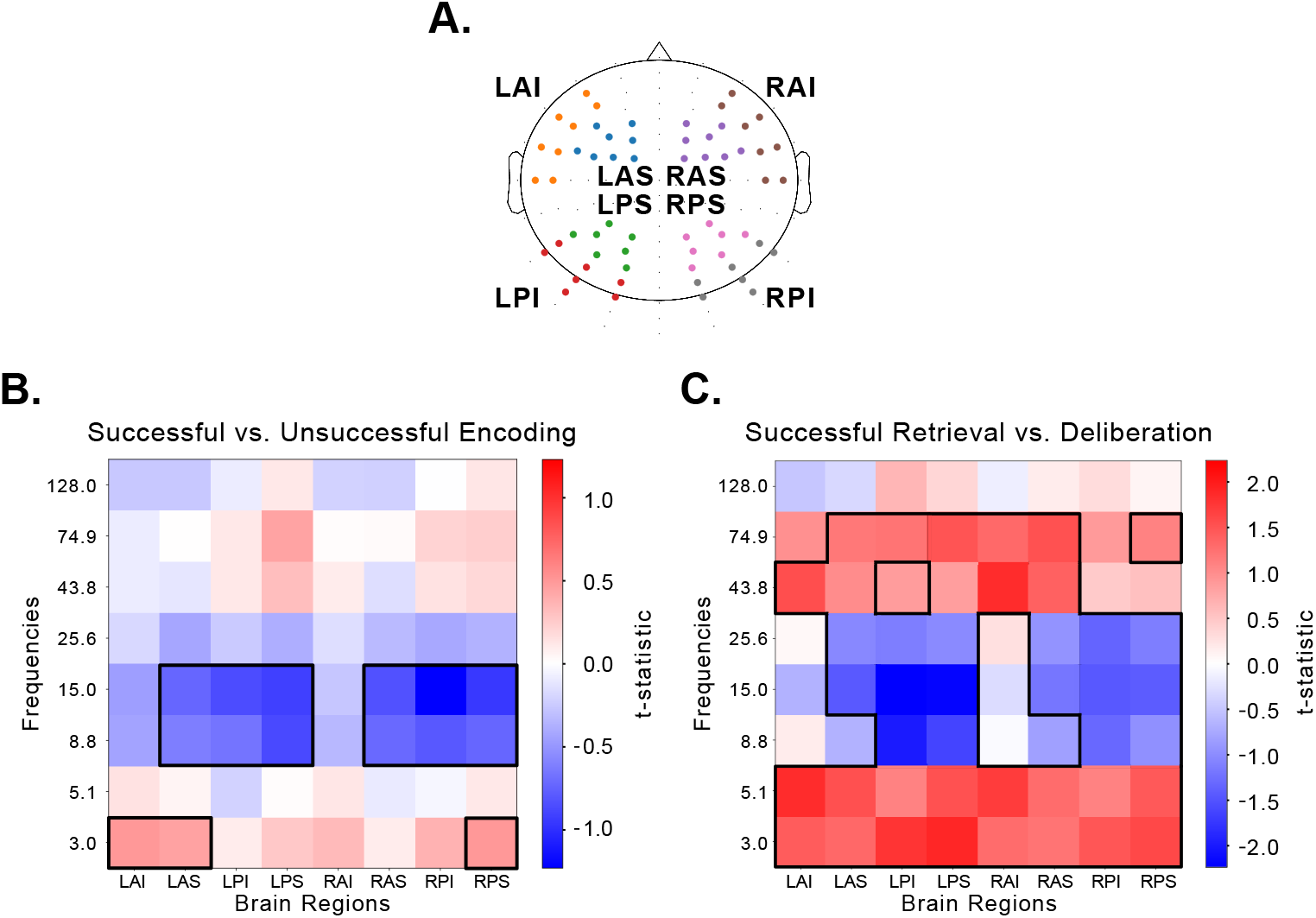
Spectral markers of successful encoding and retrieval. **A**. Diagram of ROIs on the scalp. L: left, R: right, A: anterior, P: posterior, I: inferior, S: superior. **B**. Differences in spectral power during encoding for subsequently remembered versus subsequently forgotten items. **C**. Differences in spectral power during retrieval for intervals 900ms to 50ms preceding correct recalls versus matched deliberation periods. Colors in each frequency × ROI pair correspond to the mean of independent *t*-scores for electrodes within a region of interest, averaged across subjects. Outlined frequency × ROI pairs indicate *p* < 0.05 of FDR corrected one-sample *t*-scores of the average *t*-scores within each frequency × ROI pair across subjects.

We compared average power estimates for successful recall versus matched deliberation events (see *Methods*) according to the same method. As shown in Figure 6C, successful retrieval events demonstrate similar trends to encoding, with statistically significant alpha/beta power decreases and low-theta power increases. However, the effects were much more pronounced and widespread at retrieval, with statistically significant low and high-theta increases across all ROIs as well as statistically significant gamma power increases across various ROIs in the right and left hemispheres.

Figure 7 provides a visualization of subject-level power changes at encoding (panel A) and retrieval (panel B), averaged across all electrodes. Each subject’s data appears in a separate row, with increased power (*t*-scores) shown in red and decreased power shown in blue (subjects with high recall performance appear in the upper rows). Here we see that key features of spectral *T* ^+^*A*^−^*G*^+^ appear fairly consistently across subjects at retrieval. For memory encoding, the theta increases and alpha decreases appear fairly consistently across subjects. Although encoding gamma increases appear strongly for some subjects, this effect is clearly inconsistent across our sample.

**Figure 7.**
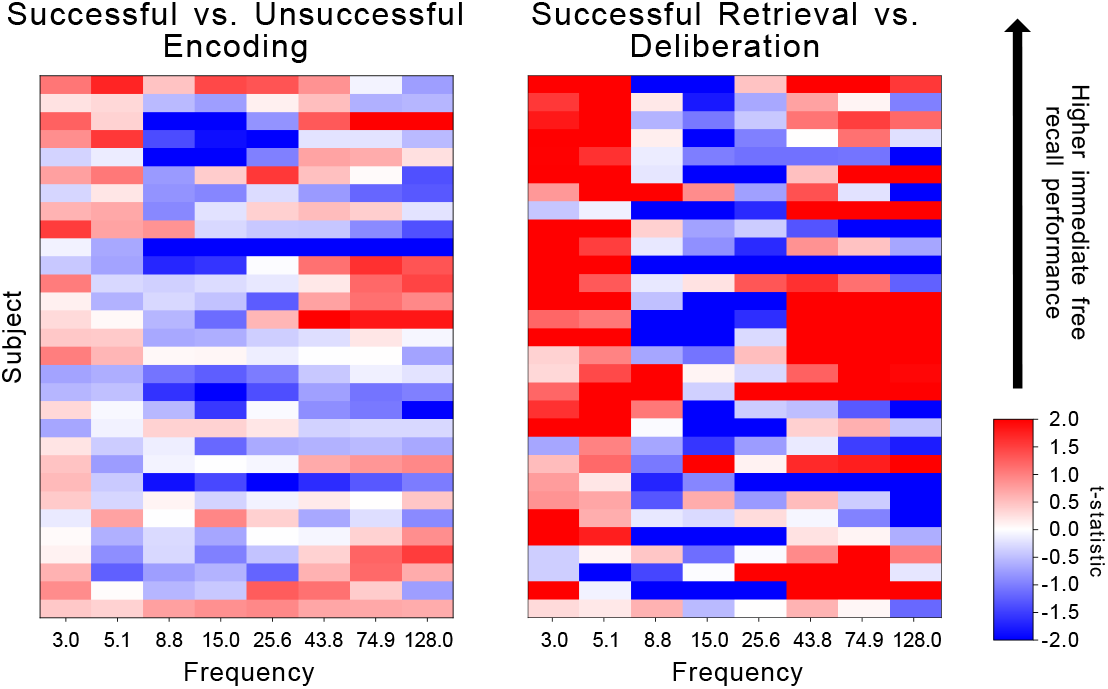
Subject-specific spectral markers of successful episodic memory in encoding and retrieval. Each row shows results from one subject, sorted by recall performance. Subject-specific independent *t*-statistics for the successful vs. unsuccessful memory encoding contrast (left) and successful retrieval vs. deliberation contrast (right) are collapsed across all electrodes. Power (t-score) increases and decreases are shown in red and blue, respectively.

These neural results generalize to those previously documented in word list tasks, with the exception of gamma power at encoding. In scalp EEG, one would expect to see broadband gamma power increases at encoding similar to those seen at retrieval. However, Sederberg et al. (2006) have indicated the possibility that these gamma-band increases at encoding are primarily driven by list position (many papers, including those from our group, have referred to gamma as high-frequency activity, HFA). Gamma appears to be particularly pronounced at the beginning of lists and trails off as lists progress, lending to the idea that gamma at encoding is more indicative of attentional orientation than a marker of successful memory (see Herweg, Solomon, & Kahana, 2020, for a discussion). Our Courier task is unique in that subjects must navigate to a business at the beginning of the encoding period prior to presentation of the first to-be-remembered item, which takes 16s on average. If gamma is in fact indicative of an initial attentional orientation, we would not expect to see strong gamma encoding effects in our Courier task.

### Multivariate analysis of mnemonic success

Whereas univariate statistics give an estimate of population-level trends for individual spectral features, they evaluate how each feature relates to mnemonic function in isolation. We leveraged multivariate pattern analyses to evaluate the joint predictive power of our entire spectral feature space. Given the strong, consistent univariate biomarkers of memory success across subjects, we predicted that multivariate classifiers could be reliably trained to decode memory states from EEG *within* a single subject, both at encoding and retrieval, and would utilize features similar to those shown in Figure 6.

We trained subject-specific multivariate classifiers to decode successful encoding and retrieval epochs on the basis of spectral features across all electrodes. We labeled encoding epochs on the basis of subsequent correct recall and retrieval epochs based on whether they preceded a successful recall or silent deliberation (see *Methods*). We used a leaveone-session-out cross-validation scheme, evaluating classifier performance as the AUC generated by the ROC function, over the predicted probabilities of events in the hold-out sessions.

Figure 8A and C summarize classifier performance across all subjects. The encoding classifiers reliably predicted memory success for data from 13 of the 29 subjects (all *p* < 0.05, permutation tests). At the group level (mean AUC = 0.53, *S E* = 0.01), the distribution of the observed AUCs was significantly greater than the theoretical null of 0.5 (*t*(29) = 4.53, *p* < 0.001, *d* = 1.19, 95% CI [0.02, 0.05]). The retrieval classifiers reliably discriminated between correct recalls and failed retrieval for all 29 subjects (all *p* < 0.05, permutation test) and at the group level (mean AUC = 0.698, *S E* = 0.01, *t*(29) = 18.99, *p* < 0.001, *d* = 4.99, 95% CI [0.18, 0.22]).

**Figure 8.**
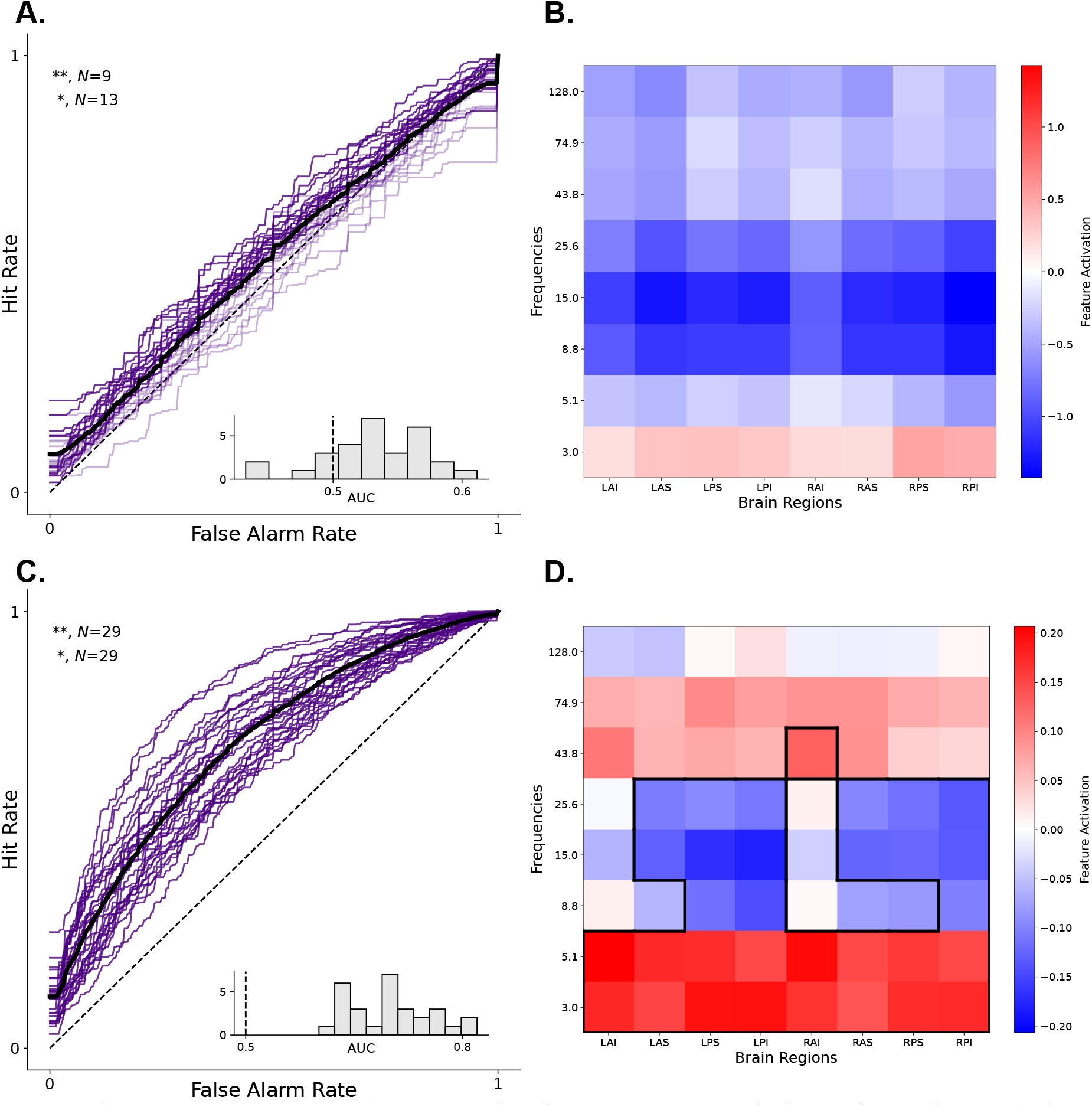
Decoding mnemonic success. Classifiers predicted memory success at both encoding and retrieval. **A**. Receiver Operating Characteristic (ROC) curves depicting the subject-level performance of the encoding classifiers. An encoding event is deemed successful if the item is subsequently recalled. Annotations in the upper left region of the ROC plots indicate the number of classifiers with significant performance (determined via permutation test). **, *p* < 0.01. *, *p* < 0.05. The lower right region of each ROC shows the distribution of observed AUCs. **B**. Feature activations for encoding classifiers, averaged across subjects. Colors in each frequency × ROI pair correspond to the mean activation value across subjects. A *t*-test against zero showed that some alpha band features were significant, but this effect did not survive FDR correction. **C**. ROCs for retrieval classifiers trained to distinguish between epochs preceding correct recalls and periods of silent deliberation (failed recall). **D**. Feature activations for retrieval classifiers, averaged across subjects. Outlined frequency × ROI pairs indicate *p* < 0.05 of FDR corrected one-sample *t*-scores of the average *t*-scores within each frequency × ROI pair across subjects.

Figure 8B and D show feature activation maps derived from the subject-specific forward-models (see *Methods* for details), with the activation patterns aggregated over electrodes in different ROIs and across subjects. In predicting whether a word will likely be subsequently recalled, the typical encoding classifier appears to rely on alpha band decreases similar to those shown in Figure 6D, but this effect was not significant after FDR correction. The feature weights for the retrieval classifiers echoed findings from our univariate contrasts for successful and unsuccessful retrieval, showing that similar spectral markers drive the predictions of these models.

### Cued and cumulative recall

We restricted our EEG analyses to the initial free recall task performed by subjects upon completion of each delivery day. Our experiment, however, also included two supplementary memory tests: A cued recall test for each delivered item (with each business serving as a cue) and a cumulative free recall test following each block of five delivery days. In cumulative free recall, subjects attempted to recall all delivered items across the five preceding delivery days (this is analogous to the classic final-free recall procedure of Craik (1970), however, since subjects performed two testing blocks separated by a 10 minute break within one session, we describe each period as cumulative recall rather than final-free recall). Because recalling an item acts as an additional learning event (Kuhn, Lohnas, & Kahana, 2018; Sheaffer & Levy, 2022), sorting encoding events based on performance of these secondary memory tasks (for EEG analyses) would be confounded by output encoding effects. Below, we report behavioral findings from cumulative free recall conditional on an item’s recall history on the prior immediate free recall and cued recall tasks.

Discrete word list tasks demonstrate two striking phenomena in cumulative free recall: long-term recency across lists and negative recency within lists. Specifically, subjects exhibit superior recall of items experienced on recent lists, but worse recall for the final items in each of those lists, reversing the positive recency effect seen with those same items in immediate recall (Craik, 1970). We therefore asked whether these phenomena would also appear in our Courier task. In addition, we wondered whether these effects would depend on whether an item was successfully retrieved in the cued recall phase. Figure 9 illustrates each of these effects. As expected, items that subjects failed to recall in both immediate tests were rarely recalled in cumulative free recall (dotted line, grey square). For items successfully recalled in both initial free and cued recall tests (solid lines and filled circles), we see a long-term recency effect, with recall rates being highest for the final delivery day and lowest for the first delivery day. For this group of items, we do not see any marked effect of within-list serial position. The subset of items that only failed immediate free recall exhibit a similar pattern with slightly lower recall. However, the subset of items that only failed cued recall exhibited a marked negative recency effect. Earlier work has established that freely recalled items from non-recency positions gain a spaced practice advantage over recency items (Craik, 1970; Kuhn et al., 2018). However, in our study, the additional learning opportunity afforded by successful cued recall provides an additional spaced learning opportunity (Allen, Mahler, & Estes, 1969; Carrier & Pashler, 1992; Soderstrom, Kerr, & Bjork, 2016). As such, it is not surprising that the negative recency effect only appeared for those initially recalled items that were not successfully recalled in response to the business cue.

**Figure 9.**
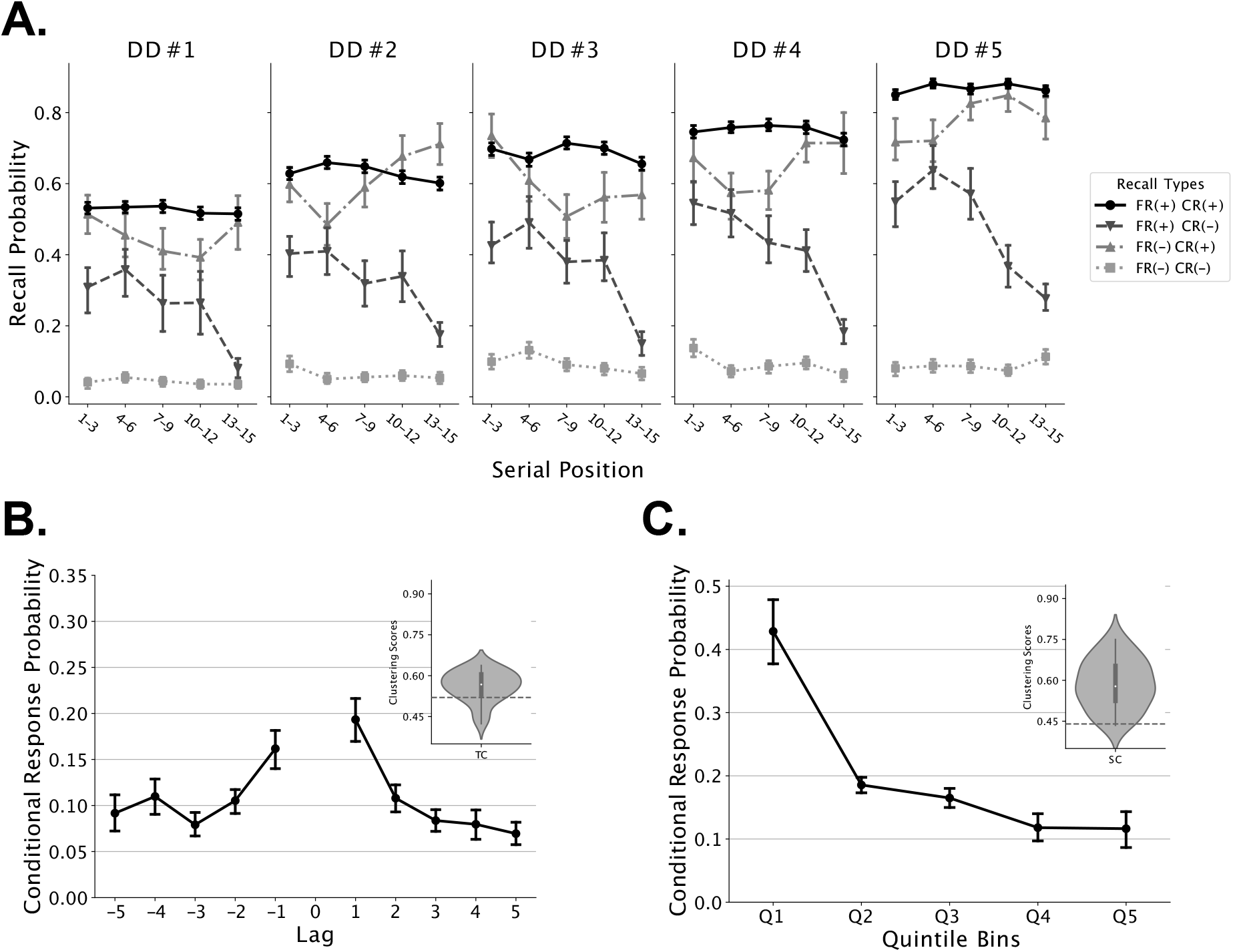
Cumulative Free Recall. **A**. Probability of recalling an item during cumulative free recall as a function of its serial position at encoding across five delivery days (DD). We categorized recall events at cumulative free recall into four conditions: FR(+/-) indicates whether subjects recalled a given item during immediate free recall, and CR(+/-) indicates whether subjects recalled a given item during cued recall. **B**. Probability of recalling an item as a function of its lag from the just-recalled item (the difference between the serial position of its predecessor and its serial position). We only included within-DD transitions in this analysis. The graph in the top right corner shows a violin plot of average temporal clustering scores during cumulative free recall. **C**. Probability of recalling an item a given spatial distance away from the just-recalled item. We included across-DD transitions in this analysis, however, we excluded same-business transitions (i.e., recalling two consecutive objects from same business location). Graph in the top right corner shows violin plot of average spatial clustering scores during cumulative free recall. Error bars indicate 95% confidence intervals, calculated using 1000 iterations of bootstrapped resampling of the data

Cumulative free recall afforded us an additional opportunity to examine the effects of temporal and spatial clustering. Because different delivery days all occurred within a single spatial environment, across-list recall transitions in this phase provided greater opportunities to observe the spatial organization of memories, while within-list transitions provided another opportunity to measure the effect of temporal organization. Figure 9B and C show that subjects exhibited robust temporal (permutation test *p* < 0.001, see *Methods*) and spatial (permutation test *p* < 0.001, see *Methods*) clustering in cumulative free recall, with comparable overall levels of clustering to those seen in immediate free recall.

## Discussion

Recent decades have witnessed enormous strides in our understanding of the neural bases of learning and memory. Using electrophysiological recordings and functional neuroimaging, cognitive neuroscientists have uncovered brain signals that predict whether an experienced item will later be remembered (Fernandez, Madore, & Wagner, in press; Hanslmayr, Staresina, & Jensen, in press), and whether a subject is about to remember a previously experienced event (Katerman et al., 2022; Herz et al., 2022). To precisely control the conditions of encoding and retrieval, researchers have predominantly studied brain signals while subjects learn and recall discrete lists of memoranda such as words or pictures – conditions that differ dramatically from recalling experienced events in our daily lives (Diamond, Abdi, & Levine, 2020)

The present study sought to examine the EEG correlates of memory encoding and retrieval in a task that embodies far greater complexity, and perhaps greater ecological validity, as compared with word list learning. In this manner, our work adds to a growing line of research embracing the study of memory in more naturalistic settings, or with greater realism, than traditional list-memory tasks (Manning et al., 2014). In the Courier task, subjects experienced items as they navigated through a complex virtual environment. Playing the role of a courier, they faced multiple “delivery days” during which they navigated to a sequence of locations (businesses), and upon arrival at each business, they learned the identity of the item that they delivered. Subjects later recalled the delivered items, either freely or in response to specific retrieval cues (see also, Herweg, Sharan, et al. (2020)). The spatial nature of our task allowed subjects to associate their experiences with a rich spatiotemporal context. By having subjects experience items only after navigating to the target business, our task created a contextual separation between learning events, more closely mimicking the way people learn in their daily lives.

Subjects exhibited striking effects of temporal and spatial clustering, recalling items in association with those experienced at nearby times and in nearby locations (see Figure 2). Temporal and spatial clustering manifested in IRTs as well, with shorter IRTs for successive recalls presented close in time and space (see Figure 3). These results lend credence to the use of naturalistic tasks to capture additional dimensions of memory organization that evade the simplicity of traditional word list tasks. As the town layout did not change across sessions, subjects learned to navigate more efficiently as they gained experience playing the game. Their improved spatial knowledge led to an increased tendency to exhibit spatial clustering of recalled items (see Figure 5D).

Spectral analyses of EEG signals revealed pronounced alpha/beta band decreases and theta band increases during both successful encoding and successful retrieval. Prior to successful retrieval, we also observed marked increases in gamma band activity. These results, along with those derived from multivariate classifier-based analyses of our data, instantiate key elements of the spectral *T* ^+^*A*^−^*G*^+^ pattern previously documented in word list tasks. Whereas retrieval data illustrated all three components of *T* ^+^*A*^−^*G*^+^ (Figure 6C), encoding data did not yield a significant gamma increase for subsequently recalled items (Figure 6B). Previous work, however, has linked the gamma increase during successful encoding to the pronounced primacy effect seen in most word list tasks (Sederberg et al., 2006; Serruya, Sederberg, & Kahana, 2014). As a result, these previous authors have hypothesized that encoding gamma increases may be indicative of an attentional boost given to early list items. The missing (or muted) gamma effect in our Courier task aligns with this hypothesis, as one would expect a reduction in primacy given the requirement to find the next business prior to each delivery (Howard & Kahana, 1999; Kahana, 2012). Taken together, our spectral analyses suggest the conservation of memory-related neural processes between traditional list-memory tasks and more quasi-naturalistic experiences.

Whereas prior research has largely studied memory for time and space in isolation, we sought to create a controlled memory paradigm to observe the joint effects of spatial and temporal organization during memory search. Our Courier task achieved this purpose, demonstrating that the order and timing of subjects’ recalls conforms to both the temporal and spatial organization of experienced events. Further, this task allowed us to determine whether neural features long-studied in traditional list-memory tasks would generalize to a more quasi-naturalistic memory paradigm. Embracing study methods and materials beyond traditional paradigms creates opportunities to capture and understand mechanisms of memory that eluded prior research. Observing a spectral *T* ^+^*A*^−^*G*^+^ of spontaneous recall, and the theta and alpha components of encoding *T* ^+^*A*^−^*G*^+^, indicates that electrophysiological studies of laboratory tasks provide a window into mechanisms of learning and recall used in our daily lives. Multivariate classifier predictions of successful memory states further support this claim. Overall, this study demonstrates the importance of spatial context in episodic memory organization, confirms the validity of naturalistic tasks in capturing neural markers of successful memory, and demonstrates predictable neural patterns of memory in scalp EEG detectable by multivariate classifiers.

## Footnotes

1 Recordings during break periods were frequently noisy as a result of these adjustments and subject movement, and were therefore excluded when calculating variances and Hurst exponents.

